# Unravelling genome organization of neopolyploid flatworm *Macrostomum lignano*

**DOI:** 10.1101/2023.04.19.537444

**Authors:** Kira S. Zadesenets, Nikita I. Ershov, Natalya P. Bondar, Nikolai B. Rubtsov

## Abstract

Whole genome duplication (WGD) is an evolutionary event resulting in a redundancy of genetic material. Different mechanisms of genome doubling through allo- or autopolyploidization could lead to distinct evolutionary trajectories of newly formed polyploids. Genome studies on such species are undoubtedly important for understanding one of the crucial stages of genome evolution. However, assembling neopolyploid appears to be a challenging task because its genome consists of two homologous (or homeologous) chromosome sets and therefore contains the extended paralogous regions with a high homology level. Post-WGD evolution of polyploids includes rediploidization, first part of which is cytogenetic diploidization led to the formation of species, whose polyploid origin might be hidden by disomic inheritance and diploid-like meiosis. Earlier we uncovered the hidden polyploid origin of free-living flatworms of the genus *Macrostomum* (*Macrostomum lignano, M. janickei*, and *M. mirumnovem*). Despite the different mechanisms for their genome doubling, cytogenetic diploidization in these species accompanied by intensive chromosomal rearrangements including chromosomes fusions. In this study, we reported unusual subgenomic organization of *M. lignano* through generation and sequencing of two new laboratory sublines of DV1 that differ only by a copy number of the large chromosome MLI1. Using non-trivial assembly-free comparative analysis of their genomes, including adapted multivariate k-mer analysis, and self-homology within the published genome assembly of *M. lignano*, we deciphered DNA sequences belonging to MLI1 and validated them by sequencing the pool of microdissected MLI1. Here we presented the uncommon mechanism of genome rediplodization of *M. lignano*, which consists in (1) presence of three subgenomes, emerged via formation of large fused chromosome and its variants, and (2) sustaining their heterozygosity through inter- and intrachromosomal rearrangements.

---

**Keywords:** *Macrostomum lignano*, whole genome duplication (WGD), autotetraploidy, subgenomic organization, chromosome rearrangements, genome heterozygosity

## Introduction

Whole genome duplication (WGD) is an evolutionary mechanism resulting in a redundancy of genetic material. WGD can create the basis for generation in future of a great phenotypic and genomic novelty (Soltis et al., 2016), and therefore it can be associated with speciation, adaptation to new environmental conditions and diversification events (Soltis et al., 2015). It is a common evolutionary mechanism in plants but rare in animals (Otto & Whitton, 2000; Mable et al., 2011; Van de Peer et al., 2017). Nevertheless, two rounds of WGD were detected in early vertebrate evolution (Holland et al., 1994; Meyer & Schartl, 1999; Dehal & Boore, 2005) that supported the predictions made by the geneticist and evolutionary biologist Susumu Ohno (Ohno, 1970).

Polyploid species can be split into two classes depending on whether the doubled genome was resulted from auto- or allopolyploidy (Garsmeur et al., 2014; Wendel et al., 2015; Van de Peer et al., 2017). Different mechanisms of genome doubling could lead to distinct evolutionary trajectories of newly formed polyploid species, many of those are allopolyploids with genomes containing two or more subgenomes derived from parental species. In case of allopolyploidy, before interspecies hybridization followed by WGD, each of the parental genomes had its own evolutionary history (Comai, 2005), own set of transposable elements (TE) and system controlling their transposition (Edger et al., 2018; Alger and Edger, 2020). TEs could be involved in intensive reorganization of duplicated genome; moreover, the differences in the TE content in parental genomes could predetermine the following evolutionary fate of the newly formed allopolyploid form and its further subgenome reorganization (Zhao et al., 2017).

A burst of TE activity (transposition, expansion and/or amplification) firstly cause changes in gene expression and following directional elimination of gene duplicates in one of the parental subgenomes making more prominent differences between them, i.e. subgenome dominance (Edger et al., 2018; Alger and Edger, 2020). Although subgenome conservation is more usual for autopolyploids (Garsmeur et al., 2014; Wang et al., 2017), some allopolyploid genomes also display genome stability and exhibit no genome fractionation, for instance the allopolyploid genome of the upland cotton *Gossypium hirsutum* (Flagel et al., 2008). In contrast to allopolyploids, autopolyploids arise through the doubling of the same or highly similar genomes also coming from individuals of polymorphic populations or subspecies. In autopolyploids, no rapid significant subgenome reorganization was usually observed (Garsmeur et al., 2014; Wang et al., 2017) and the most of gene duplicates tend to be retained (Zhao et al., 2017), gene duplicates could differentiate not in the form of synteny blocks or subgenomes, but as individual gene pairs (Zhao et al., 2017).

Genome studies on species underwent a recent WGD are undoubtedly important for understanding one of the crucial stages of genome evolution. At the same time, these studies are rather complicated with newly formed polyploid genomes that are deciphered in two homologous (or homeologous) chromosome sets and therefore contain extended paralogous regions showing a high homology level. In result, assembling polyploid genomes often appears to be a challenging task due to the difficulties in distinguishing between paralogous sequences of genome. In case of autopolyploids, additional problem could derive from nearly identical haplotype variants in paralogous chromosome regions. Assembling up to chromosome level could be complicated also by structural rearrangements of chromosomes, such as indels, duplications, translocations with breakpoint regions enriched for repeats (Ming and Wai, 2015; Zhang et al., 2020; Zadesenets & Rubtsov, 2018). Not surprisingly, assemblies of polyploid genome can contain many collapsed or ‘mosaic’ (chimeric) sequences, making almost impossible the comprehensive analysis of polyploid genome organization and recovering of its evolutionary history from a WGD event to the modern state. Post-WGD evolution of polyploid species usually includes rediploidization stage, first part of which could be cytogenetic diploidization. This process results in the formation of species, whose polyploid origin might be hidden by disomic inheritance and diploid-like meiosis (Svačina et al., 2020). These species can be recognized as polyploids with difficulty.

Numerous studies on genome evolution were conducted mainly on plants (Spoelhof et al., 2017), whereas only few species with a recent WGD event in their evolutionary history were involved in genome studies in animals (Session et al., 2016). One group of such species includes free-living marine flatworms belonging to the genus *Macrostomum*, namely *M. lignano, M. janickei*, and *M. mirumnovem* (Zadesenets et al., 2016, 2017a,b, 2020; Zadesenets & Rubtsov 2018, 2021). Furthermore, they belong to two phylogenetic lineages (Schärer et al., 2020) with two independent WGD events and similar karyotypes’ reorganization resulted in diploid-like karyotypes (Zadesenets et al., 2020). The revealed difference between karyotypes of *M. lignano* and *M. janickei* belonging to the same lineage was limited to the doubling of the copies of the largest chromosome in the *M. janickei* karyotype and the absence of the 28S rDNA cluster in its large chromosomes (Zadesenets et al., 2016, 2017a,b). Although in both *M. lignano*/*M. janickei* and *M. mirumnovem* lineages WGD events occurred at roughly the same time, the large scale changes in the following karyotypic evolution were different. Evolution of their karyotypes included chromosome fusions with following formation of large metacentric chromosome(s). In *M. mirumnovem*, chromosome rearrangements accompanied with the expansion and amplification of repetitive sequences, formation of B chromosomes, and prominent karyotype instability linked with structural and numerical chromosome variation. Altogether, this suggested a massive genome reshuffling after a recent WGD probably included interspecies hybridization (Zadesenets et al., 2020, 2021). In contrast to *M. mirumnovem*, the karyotypes of *M. lignano* and *M. janickei* are more stable and their diversity is mainly associated with varied copy number of the large chromosome (Zadesenets et al., 2016, 2020). The aneuploid frequency in these species was low in natural populations but increased in the laboratory cultures (Zadesenets et al., 2016, 2020). Remarkably, in both species the largest chromosome consists of the extended regions paralogous to the regions of remaining unfused small metacentrics (Zadesenets et al., 2017a,b). In *M. lignano,* a few small deletions and inversions distinguishing paralogous regions of the large and small chromosomes were described (Zadesenets et al., 2017b). Based on the results of molecular cytogenetic studies, we suggested the possible origin of *M. janickei* from *M. lignano* by doubling the largest ancestral chromosome.

For further studies on genome evolution in the genus *Macrostomum*, the genome assemblies of the *Macrostomum* species are required. A high karyotype diversity and pronounced genome instability in *M. mirumnovem* provides serious difficulties for its genome sequencing. Earlier a few attempts were made to decipher the genome of *M. lignano* (Wasik et al., 2015; Wudarski et al., 2017), but the resulting assemblies suffered from substantial fragmentation despite PacBio deep sequencing that was applied: N50 of ML2 and Mlig_3_7 assemblies were 64 Kb and 246 Kb, respectively. Furthermore, the Mlig_3_7 assembly appeared to be ∼700 Mbp in size, while the size of gDNA in cell nuclei was estimated as 500 Mbp (Wudarski et al., 2017). This does indicate that despite the relatively small size of the *M. lignano* genome, its assembling appeared to be a challenging task. The authors suggested that the assembly fragmentation was resulted from an unusually high genome repetitiveness providing by both chromosomal duplications and pronounced expansion of simple DNA repeats and transposons (Wasik et al., 2015). Thus, routine approaches to assembling the neopolyploid *M. lignano* genome yielded no satisfactory results. Clarification of unexpected complexity of the *M. lignano* genome required a new approach to its analysis.

Encouraged by these findings, we applied the original analysis of self-homology within the reference genome assembly of *M. lignano*, combining with the comparative analysis of WGS data of microdissected library of the *M. lignano* largest chromosome (MLI1) and the two newly established laboratory sublines with two and four copies of the MLI1, respectively. Here we reported subgenomic organization of the *M. lignano* genome and discussed the possible mechanisms of genome evolution in the genus *Macrostomum*.

## Results

### Self-homology within the reference genome assembly of M. lignano

Based on our previous studies, we inferred that *M. lignano* is a neotetraploid species (Zadesenets et al., 2017a,b), and therefore its genome mainly consists of duplicated DNA sequences, with some of the duplications being lost or altered due to rediploidization. For detailed analysis of genome organization, we performed a copy-number analysis of the earlier published genome assembly, Mlig_3_7 (Wudarski et al., 2017). This genome assembly like others (Wasik et al., 2015) was generated on the sequencing data obtained for the inbred line DV1. We identified paralogous DNA sequences by looking for long-range self-homologies between the scaffolds and slicing the entire assembly onto PBs. Each PB comprised a target genome fragment together with its fully aligned homologs (if any were found), so that the copy number of a target was unambiguously determined (Fig. S1). Unexpectedly, the performed analysis showed the prevalence of PBs containing three copies of paralogous DNA fragments across the genome assembly Mlig_3_7 (Fig. 1A). Actually, these three-copy PBs accounted for about 69.6% of the assembly length, whereas 2.2%, 14.7%, and 11.9% of genome were presented by unique, duplicated and four-copy DNA sequences, respectively. The remaining 1.6% of the genome was presented by multicopy PBs. We should note that these estimates are based solely on analysis of the draft genome assembly, which is certainly error-prone. Moreover, incorrect copy-number representation arising from improperly collapsed or expanded haplovariants is a quite common error in genome assembling of polyploid species (Yang et al., 2017; Zhang et al., 2020).

**Figure 1.**
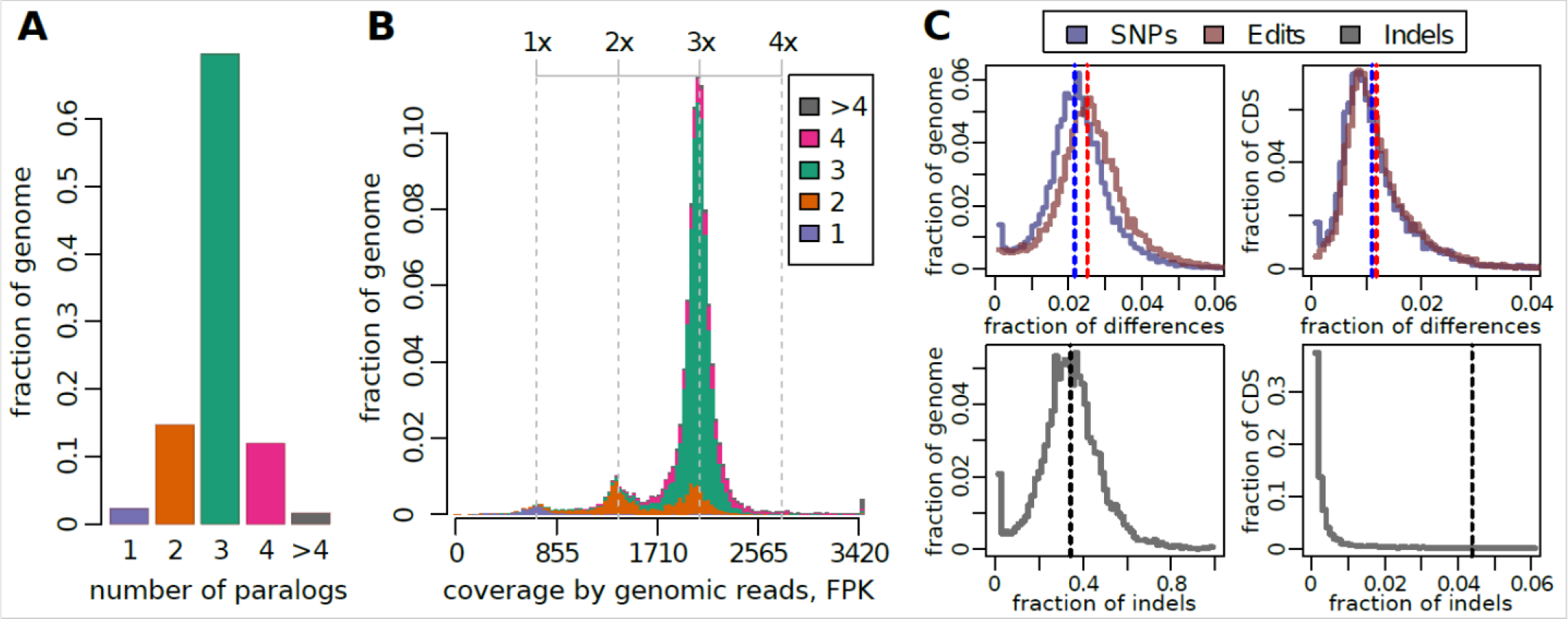
Copy number variation and divergence of paralogous sequences in the reference genome assembly *M. lignano*, Mlig_3_7. (A) Proportion of unique and multi-copy sequences in the Mlig_3_7 genome assembly, based on grouping of chained pairwise self-alignments into paralogous blocks (PBs). (B) Distribution of aggregated coverage of PBs by short-read genomic library (fragments per kilobase of locus length, FPK). For each PB, the sum of individual coverage values of all its members was used. The resulting data are grouped by the number of members in PBs (color scheme is the same as in panel A). It is seen that many two-copy and four-copy PBs actually have three-fold read coverage (upper axis), indicating incomplete or excessive representation of paralogous regions in the Mlig_3_7 assembly. (C) The distribution of PBs by SNP rate, Levenstein edit distance and indel rate computed over the whole sequences or just their protein-coding fraction (CDS). Dashed lines denote the total frequences computed for the entire genome.

To estimate the contribution of this type of errors, we evaluated copy number of PBs based on their total coverage by short reads of the gDNA library that was originally used for the assembly. We suppose that the discrepancy between the copy number of PB and the multiplicity of its short-read coverage would indicate a copy number error in the assembly. As a result, about 85% of the genome was indeed found in three copies (Fig. 1B). Unique, duplicated and four-copy sequences comprised 2.4%, 9.3%, and 1.7% of the genome assembly, respectively. The remaining 1.6% of the genome was presented by multi-copy sequences. Most of the four-copy PBs and about half of the two-copy PBs actually have a threefold coverage by raw reads, that is, they most likely contain erroneously represented paralogs.

To sum it up, the genome of worms from the inbred line DV1 consists of three paralogous elements. On a large scale (at least 5kb), loss of the triploid state was estimated to cover about 12% of the genome, yet without taking into account differences within aligned paralogous sequences. As for the latter, we also assessed the divergence of the identified paralogs from their chained pairwise alignments (Fig. 1C). The estimated SNP rate was 2.2% over the entire genome and 1.1% over its protein-coding sequences, supporting the suggestion about a recent WGD in the genome evolution of *M. lignano*. It is harder to interpret indels from chained alignments, since in fact they may also represent inversions and translocations that are not accompanied by the actual loss of sequences. Considering SNPs and inserts of any length as single base edits (Levenshtein distance), the resulting divergence of the paralogous loci slightly increased and reached 2.5%. However, if we take into account the length of indels, then the differences reach 36% for the entire set of aligned paralogs and 5.5% for its protein-coding sequences. It is noteworthy that the distributions of all estimates of differences for individual PBs are unimodal (Fig. 1C), which indicates fairly similar divergence rates for all three paralogs. Taken together, while the observed large-scale differences between all three paralogous copies are pronounced (about 50% identity), they are mainly presented by indels, while the low-scale homology (< 1 kb) of paralogs remains quite high (98%). As a consequence 93% of the read pairs from genomic library were mapped to multiple loci in the reference genome assembly.

Taking into account that worms from the highly inbred line DV1 were used for sequencing of the *M. lignano* genome (Wasik et al., 2015; Wudarski et al., 2017) and that the euploid karyotype of *M. lignano* was suggested to be a hidden tetraploid (Zadesenets et al., 2017a, b), the finding three large paralogous genome parts in the haploid genome assembly Mlig_3_7 does raise the question of how these subgenomes could be mapped to chromosomes of the euploid worms characterized with a hidden karyotypic tetraploidy. To address and solve these questions, we established two sublines of the DV1 line differed with the copy number of the MLI1 that covered the most part of the genome assembly Mlig_3_7. By comparing genomes with different copy number of the largest chromosome, we hoped to differentiate paralogous sequences belonging separately to small and largest chromosomes.

### The karyotype variability in the inbred line DV1 and establishing its derivatives DV1_8 and DV1_10

Karyotyping of worms from the original inbred DV1 line revealed a high frequency of aneuploid specimnes (Zadesenets et al., 2016, 2020). The observed aneuploidy was asssociated mainly with one or two additional copies of the chromosome MLI1. The high frequency of tri- and tetrasomics on the MLI1 provided us possibility for deciphering DNA content of this chromosome by comparative genomic analysis of specimens having species-specific *M. lignano* karyotype and karyotype with two additional copies of the MLI1.

To collect worms with the required karyotype variants 2n=8 and 2n=10 for establishing the DV1_8 and DV1_10 sublines, we karyotyped 50 specimens from the DV1 line. Karyotyping revealed seven worms with the normal karyotype 2n=8 (14%), 28 worms with trisomy on the MLI1, 2n=9 (56%), 14 worms with tetrasomy on the MLI1, 2n=10 (28%), and one specimen with an extra small chromosome additional to the normal *M. lignano* karyotype, 2n=9 (2%). The extra small chromosome was smaller than the smallest A chromosome and was referred to B chromosomes (Zadesenets and Rubtsov, 2021). Each of the sublines, DV1_8 and DV1_10, was established from a pair of worms.

Among the karyotyped worms-founders for pools DV1_8A-E and DV1_10A-E, the only founder (the DV1_8B pool) showed the 2n=12 karyotype distinct from the expected 2n=8. Besides an additional copy of the MLI1, it has three additional small chromosomes (Fig. S2). For estimation of worm viability and reproduction level, we counted in each pool the total number of F1 offsprings, number of died worms and worms showing any morphological or behavioral abnormalities (Table 1). The comparison between the DV1_8A-E and DV1_10A-E pools showed that the DV1_8A-E were characterized with the higher total number of offsprings (N), but also with the higher level of mortality and morphological abnormalities than the DV1_10A-E (Table 1).

**Table 1.** The assessment of offspring (N) of the generation F1 for pools DV1_8A-E and DV1_10A-E.

| Subline | ID pool | Parental worms | Karyotype | Number worms among offspring F1 |  |  |  |
| --- | --- | --- | --- | --- | --- | --- | --- |
|  |  |  |  | Total number | Normal<sup>†</sup> | Died<sup>‡</sup> | Abnormal<sup>§</sup> |
| DV1_8 | A | 20 worms | 2n=8 (100%) | 228 | 196 | 11 | 20/1 |
|  | B | 20 worms | 2n=8 (95%)<br>2n=12 (5%) | 128 | 88 | 15 | 23/2 |
|  | C | 20 worms | 2n=8 (100%) | 214 | 184 | 11 | 18/1 |
|  | D | 20 worms | 2n=8 (100%) | 171 | 139 | 24 | 7/1 |
|  | E | 20 worms | 2n=8 (100%) | 210 | 180 | 15 | 12/3 |
|  |  |  | Total | 951 | 787<br>(82.75%) | 76 (7.99%) | 80/8<br>(8.41%/0.84%) |
| DV1_10 | A | 20 worms | 2n=10 (100%) | 204 | 194 | 4 | 6/0 |
|  | B | 20 worms | 2n=10 (100%) | 140 | 135 | 4 | 1/0 |
|  | C | 20 worms | 2n=10 (100%) | 193 | 184 | 5 | 2/2 |
|  | D | 20 worms | 2n=10 (100%) | 125 | 121 | 3 | 1/0 |
|  | E | 20 worms | 2n=10 (100%) | 193 | 185 | 7 | 1/0 |
|  |  |  | Total | 855 | 819<br>(95.79%) | 23 (2.69%) | 11/2<br>(1.29%/0.23%) |

<sup>†</sup>alive worms with normal morphology and behavior; <sup>‡</sup>died worms; <sup>§</sup>worms of small size/ worms with other abnormalities. In 20 days after isolation of hatchlings, we counted worms showing body sizes smaller than size of normal adult worms. They were called ‘small worms’.

Based on the number of hatchlings, viability, morphology and behavior of the worms, three pools for each of the sublines DV1_8 and DV1_10 (DV1_8A, DV1_8C, DV1_8E, DV1_10A, DV1_10C, DV1_10E, Table 1) were selected for further experiments.

### Assembly-free estimation of ploidy and retained haterozygosity in genomes of the DV1_8 and DV1_10 sublines

Sequencing data for six DNA libraries obtained from DV1_10A, C, E worm pools showed high reproducibility of the genome coverage profile (Fig. S5). The same reproducibility was observed in genome coverage profiles of five DNA libraries from the DV1_8A, C, E worm pools. The only replicate (one of two libraries from the DV1_8A pool) was characterized by noticeable deviation from the average profile, which was found to be attributed to positive GC bias introduced most likely at the step of DNA library amplification. We merged all replicates from the DV1_10A, C, E worm pools and five replicates from the DV1_8A, C, E worm pools to increase the overall sequencing depth.

The obtained NGS data were inappropriate for *de novo* assembly of polyploid genome, but they appeared to be sufficient to conduct a comparative copy-number analysis and evaluate the level of the retained genome heterozygosity. We took into account that the reference genome assembly Mlig_3_7 contains a number of errors, which should adversely affect the reliability of the copy-number analysis. Therefore, we performed the comparative reference-free analysis of the DV1_8 and DV1_10 libraries based on k-mer frequency spectra. Despite of its limitations, this type of analysis allows accurate estimation of major genome properties such as size, heterozygosity, repetitiveness, and even ploidy, based solely on raw sequencing data (Ranallo-Benavidez et al., 2020).

It was revealed that the DV1_8 and DV1_10 genomes share over 99% of 35-mers (i.e. unique subsequences of length 35), meaning that there is no pronounced subline-specific changes in their sequence content, except for the changes in copy number of sequences. This circumstance allowed us to represent the distribution of k-mer frequencies in 3D space (Fig. 2), with marginal distributions corresponding to the frequency spectra of each subline. Each k-mer spectrum is composed of a series of equidistant peaks that correspond to the frequency of the underlying sequences within a genome. In this way, the area of the lowest peak corresponds to the proportion of unique haplotype-specific sequences, while the highest peak refers to completely homozygous sequences common to all haplovariants. In the 3D space defined by DV1_8 - DV1_10 pair, we observed five major peaks of frequencies, which, when expressed as fold coverage of the haploid genome, were as follows: (×1; ×2), (×2; ×2), (×2; ×4), (×3; ×4), and (×4; ×6) (Fig. 2).

**Figure 2.**
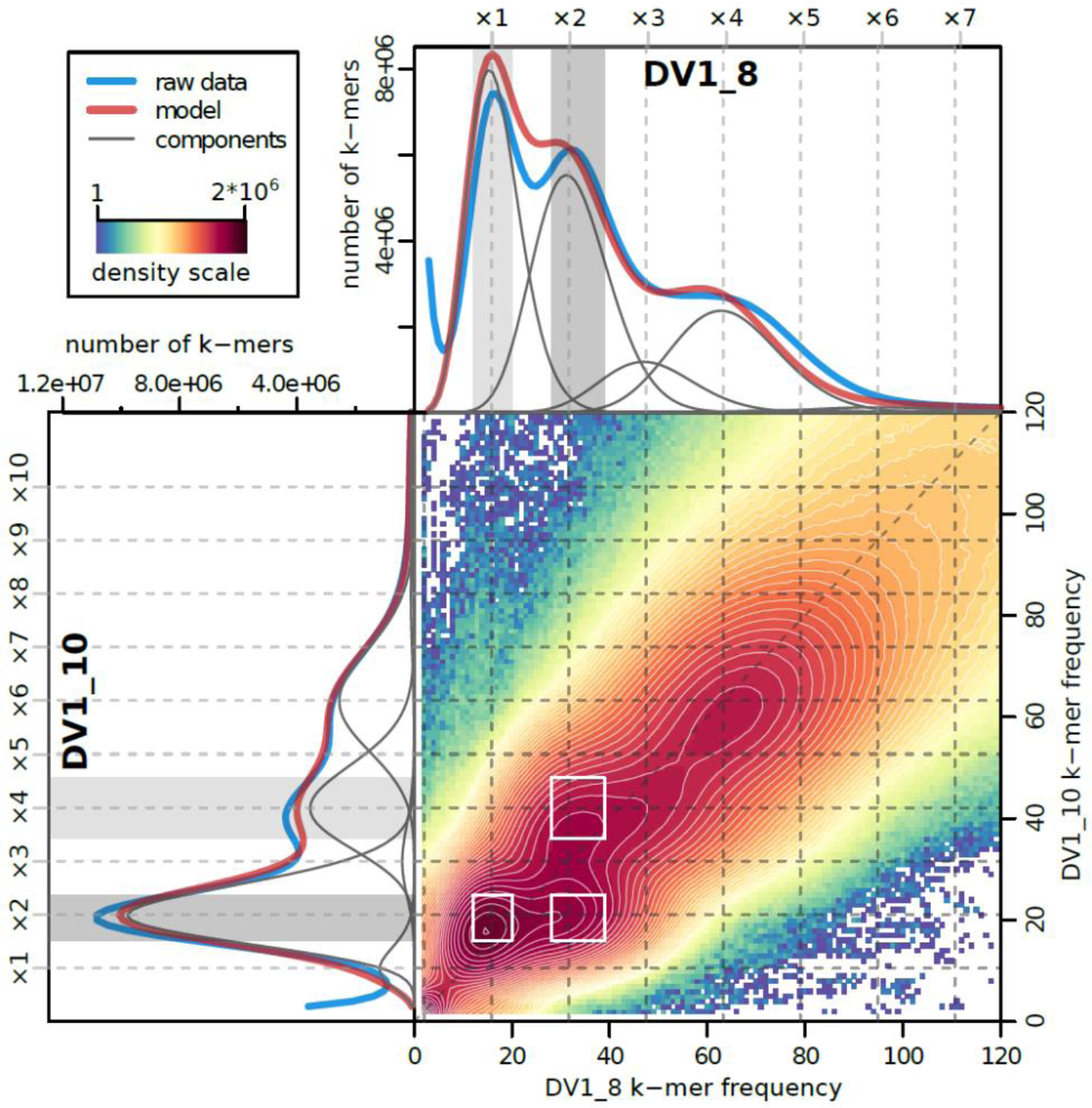
Spectra of k-mer frequencies for the sequenced DV1_8 and DV1_10 datasets. Density plot in the bottom right corner represents the 3D frequency spectrum of 35-mers that are shared between the DV1_8 and DV1_10 datasets. Density color scale depicts the number of k-mers having given frequencies. The supplementary histograms along its axes (top and left sections) show the corresponding marginal k-mer distributions for either DV1_8 or DV1_10 libraries (thick blue line), fitted mixed negative binomial model (thick red line) and its components (thin grey lines). Top and left scales and dashed grey lines show the coverage multiplicity factor corresponding to the copy number of genomic loci. Three subsets of subgenome-specific k-mers that were extracted and used for classification of the genomic scaffolds (S and L subgenomes), are depicted by white rectangles at 2D plot as well as by grey areas at 1D histograms.

Given the highly inbred state of the sublines DV1_8 and DV1_10 and their chromosome sets, the simplest genome configuration that fits this distribution required a set of three subgenomes (haplovariants) denoted as S, L_1_, and L_2_. Under this scenario, the DV1_8 genome is composed of one diploid subgenome and two haploid subgenomes (SSL_1_L_2_), while the DV1_10 genome was transformed into hexaploid form SSL_1_L_1_L_2_L_2_ (Table S2).

To further study of the heterogeneity and heterozygosity level of these three subgenomes, we fit the GenomeScope2.0 polyploid model (Ranallo-Benavidez et al., 2020) modified in several aspects to fit to our data and suggested assumptions (Table S1) (for details see the section “Material and Methods”). To keep the analysis comprehensive we intentionally left the estimators for heterozygosity between two homologous copies of each haplovariant ignoring the fact of the high inbreeding level in the DV1 line.

As a result of the model fit, only three of the eight estimated heterozygosity rates were substantially different from zero (Table S1). More specifically, the predicted contribution of heterozygosity between copies of each subgenome to the total variation was negligibly small, which is in a good agreement with the highly inbred state of the sublines. The overall SNP variation in the *M. lignano* genome appeared to be 2.6%. The inferred divergence between L_1_ and L_2_ subgenomes was 1.39%, while 1.58% referred to dissimilarity between S and any of L_1_ and L_2_ haplovariants. It is noteworthy that the applied poset-based model mostly capture small-scale differences such as SNPs, so the obtained estimates of divergence are close to the values of SNP rate we obtained from self-alignment of the genome assembly Mlig_3_7. The fact is that the sensitivity of k-mer-based approaches is directly limited by the chosen k-mer size, and estimating the actual contribution of larger structural variants, especially those involving DNA repeats, remains beyond the capabilities of the GenomeScope model. At the same time, large rearrangements, including inversions and translocations, can effectively promote genetic isolation by limiting meiotic recombination. In this respect, using the FISH technique we have previously shown, that the order of loci within the large chromosome is altered by inversions compared to that in small metacentrics (Zadesenets et al., 2017b). Nevertheless, the conducted reference-free analysis further supported the existence of three genetically isolated subgenomes that formed genomes of both DV1_8 and DV1_10 inbred sublines.

### Determination and validation of subgenome-specific sequences in the Mlig_3_7 genome assembly

According to the karyotyping data, the worm pools from the DV1_8 and DV1_10 sublines differed only by doubling the copies of the MLI1. At the same time, the analysis of sequencing data showed that worms from the DV1_8 subline showed an unexpected type of polyploidy (SSL_1_L_2_), while the worms from the DV1_10 subline exhibited a balanced hexaploid genome (SSL_1_L_1_L_2_L_2_). In this regard, the copies of the MLI1 should differ and correspond to the subgenomes L_1_ and L_2_. Consequently, the relative read coverage of these subgenomes in the DV1_10 subline should be twice a high as in the DV1_8, which makes it possible to divide the reference assembly into two sets belonging to small and large chromosomes, respectively.

In concordance with our expectations, the bivariate distribution of read coverage computed across 5-kb genomic windows for the two data sets showed two major density peaks corresponding to S and L_1_/L_2_ subgenomes (Fig. 3A). However, the distribution was distorted by several kinds of bias. First, clusters of under- and over-covered regions stretched in three diagonal directions are caused by incorrectly expanded or collapsed allelic variants in the reference assembly. It is noteworthy that this sort of bias was most relevant to the pair of subgenomes L_1_ and L_2_ (the leftmost diagonal). There is also a mapping bias that results in a shortened distance between the two major peaks along the DV1_8 axis (1.8-fold difference instead of the expected 2.0-fold). It is driven by a large proportion of multi-mapped reads that are equally similar to at least two subgenomes. The probability of their placement to any of subgenomes in the reference assembly (SL_1_L_2_) is 1/3, while in DV1_8 (SSL_1_L_2_) the probability of their origin from the S variant is 1/2, and from the each of L_1_ or L_2_ variants is 1/4. We have shown that this bias disappears if all multi-mappable genomic positions have been excluded from consideration (Fig. S6). Finally, the increased spacing between the peaks along the DV1_10 axis is caused by another unidentified source, which is also present in the k-mer spectra of raw sequencing data (Fig. 2).

**Figure 3.**
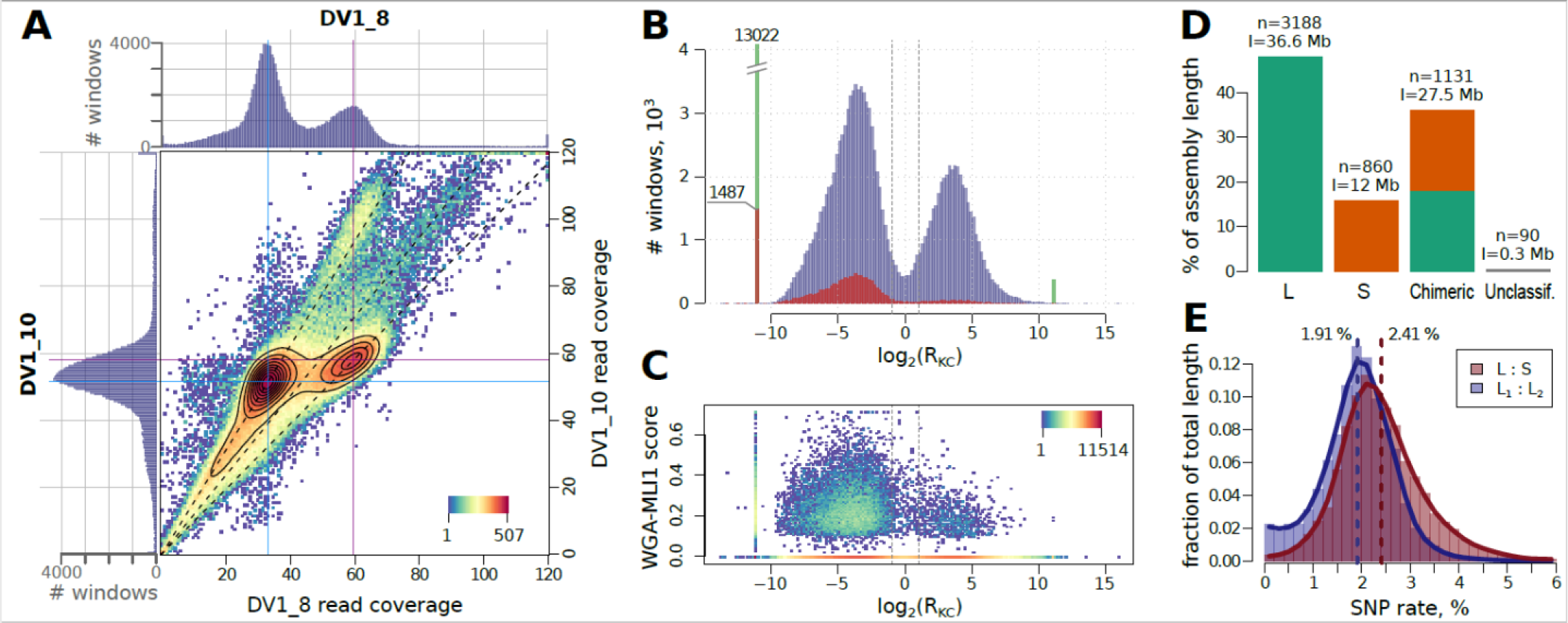
Discrimination of subgenome-specific loci in the Mlig_3_7 assembly using the NGS data for the DV1_8 and DV1_10 sublines. (A) 2D density plot of the genome coverage distributions of the DV1_8 and DV1_10 libraries, computed over 5-kb sliding windows (the color scale denotes the density of the windows). Top and left histograms illustrate coverage distributions of each library. Blue and red lines mark summits of two main peaks of the coverage. Dashed black lines show the axes of bias arising from unresolved or overrepresented paralogous pairs in the assembly. (B) Bimodal distribution of the log_2_(R_KC_) (log_2_-ratio of S to L k-mer counts) metric computed over the same 5-kb genomic windows. Spiked green bars are made of windows with infinite log_2_(R_KC_) due to zero counts of either S- or L-specific k-mers. Dashed lines show the threshold of two-fold difference that was used for subsequent classification of windows into S and L classes. Red-colored bars show the fraction of windows having positive signal from MLI1-specific WGA-seq library. Density plot (C) shows the distribution of this MLI1-specific signal along the values of log_2_(R_KC_) in the corresponding windows. (D) Proportion of determined subgenomic classes in the assembly Mlig_3_7. The captions above the bars are the number of scaffolds (n) and their total length (l). (E) Distribution of SNP frequency in either L1-L2 or L-S pairs of paralogs taken from the classified three-copy “S-L1-L2” PBs. Dashed lines show the frequences computed for the whole set of “S-L1-L2” PBs.

In view of the foregoing we decided to come back to k-mer analysis to refer k-mer spectra to retriek-mers belonging to the S or to the both L subgenomes, and as well as belonging to divergent alleles of L_1_ and L_2_ variants. Of five peaks in the (DV1_8; DV1_10) frequency spectrum, the peak (×1; ×2) correspond to k-mers, specific to either L_1_ or L_2_ subgenomes, and the peak (×2; ×4) contains k-mers, specific to both L subgenomes, i.e. to both different copies of the largest chromosome (Fig. 2, Table S3). On the other hand, k-mers that are specific to a set of small chromosomes (subgenome S) fall into the peak (×2; ×2). The extracted subsets (marked on Figure 3) were mapped to the genome assembly and used to calculate genome coverage over 5 kb windows. To assign the windows to the S and L subgenomes, we took the log_2_-ratio of normalized coverage by S-specific k-mers to those by L-specific k-mers (log_2_(R_KC_)). As shown in Figure 3B, the number of windows formed a bimodal distribution with peaks equidistant from zero. At the two-fold difference threshold, 32% and 64% of the windows were assigned to the S and L subgenomes, respectively, which is consistent with the expected subgenome proportions.

To confirm that the S, L_1_ and L_2_ subgenomes correspond to sets of small and different copies of the large chromosome, we mapped the sequenced microdissected DNA library derived from the MLI1 (WGA-MLI1) and computed the breadth of coverage for the same 5-kb genomic windows that were already split into two classes S and L. As shown on Figure 3B,C, the windows from class L (those with negative log_2_(R_KC_) values) mostly coincide with the high values of the WGA-MLI1 coverage. It should be noted that the WGA-MLI1 DNA library is characterized by the predominance of multi-mapped reads and extremely uneven coverage of the genome, amounting to about 1% of its length. Therefore, many windows in class L that did not contain uniquely mapped WGA-MLI1 reads had a coverage value of zero. We should note that a portion of S-class windows had an unexpectedly high coverage with the WGA-MLI1 library, which is likely attributed to the already mentioned inaccuracies in the genome assembly and short-read mapping. Nevertheless, the WGA-MLI1 data provided evidence that the L_1_ and L_2_ subgenomes correspond to differing copies of the largest chromosome MLI1 in the karyotypes of the DV1_8 and DV1_10 sublines.

Based on the classified genomic windows, we were able to classify the assembly scaffolds and assess the level of chimerism between the S and L subgenomes in the reference assembly (Fig. 3D). At the chosen thresholds, only 90 of 5269 scaffolds remained unclassified due to the insufficient k-mer coverage. It was found that 1131 scaffolds, which make up 36% of the assembly length, are chimeric. Finally, 3188 and 860 scaffolds were assigned to the L and S subgenomes, respectively. However, we cannot rule out or take into account additional chimerism within the L class since the performed analysis did not distinguish the L_1_ and L_2_ subgenomes.

We applied the same classification scheme to the previously defined PBs and selected those containing three non-chimeric paralogs – one S and two L members. This strictly selected set of paralogs was used to compare the divergence between the three subgenomes (Fig. 3E). SNPs are 1.3 times more frequent in the pair L-S than in the pair L_1_-L_2_ (2.41% versus 1.91%). The same ratio was derived from the initial reference-free analysis of k-mer spectra. When it takes long insertions into account, the overall divergence rate is estimated at 36.46% in L-S pair and 32.08% in L_1_-L_2_ pair.

Since divergence of homoeologs can be associated with asymmetric expansion of TEs, we also explored whether there are substantial differences between S and L_1_L_2_ subgenomes in terms of repetitive content. However, no significant differences in proportions of predicted classes or families of DNA repeats were observed (Fig. S7).

Taken together, the results support the existence of three homeologous subgenomes within both DV1_8 and DV1_10 genomes, two of which correspond to the variants of the largest chromosome MLI1, and the third corresponds to a set of three small chromosomes, MLI2-MLI4.

## Discussion

Earlier we described karyotypic rediploidization after recent WGD events in some species of the genus *Macrostomum*. In *M. lignano*, it included the fusion of one set of ancestral chromosomes into one large metacentric chromosome with following retention of the extended paralogous chromosome regions. However, the question about the genomic rediploidization in this species remained open. Despite the relatively small size of the *M. lignano* genome and application of long read technique for its sequencing, the quality of the genome assembly appeared to be not completed. One of the possible reasons for that could be a high content of simple DNA repeats and transposon elements in the *M. lignano* genome (∼75% of genome sequence) (Wasik et al. 2015). Additional complication could derive from a high homology of DNA in paralogous chromosome regions. In this study, we unexpectedly revealed DNA heterozygosity in copies of the largest chromosome in *M. lignano* (DV1 line). Given the inbred state of the DV1 line, we suggested genetic isolation of the largest chromosome copies or possible heterozygosity generation by TEs. The latter can facilitate genetic variation due to increasing mutation rate. It was proposed that TEs can maintain genome-wide heterozygosity and may provide a basis for evolution under inbreeding (De Kort et al., 2022). The *M. lignano* genome indeed contains many simple DNA repeats and TEs, but they showed uniform distribution across the *M. lignano* genome, though in inbred line we revealed heterozygosity only in the largest chromosome. Based on these data, genome of *M. lignano* might be considered as combination of three genetically isolated subgenomes; one, diploid, associated with small metacentrics, and two others, haploid, associated with different copies of the largest chromosome MLI1. Such genome organization could explain also, why the haploid assembly Mlig_3_7 is about 1.5 times larger than the genome size estimated with flow cytometry (Wudarski et al., 2017).

Autopolyploids possess a genomes having doubled set of homologous chromosomes, resulting in polysomic inheritance (Spoelhof et al., 2017). Apparently, the formation of the chromosome MLI1 with its rapid structural alterations led to genetic isolation and further independent evolution of largest chromosome and small metacentrics, resulting in disomic inheritance. We suggested that formation of the MLI1 through chromosome fusions followed by extensive rearrangements could be uncommon mechanism to sustain heterozygosity in subgenomes deciphered in differing copies of the chromosomes MLI1.

As far as we know, the uncommon organization of the *M. lignano* genome is described by us for the first time. However, fusion of all ancestral chromosomes into one large chromosome was described earlier in parthenogenetic nematode species, *Diploscapter pachys* (Fradin et al., 2017). The unichromosomal karyotype of *D. pachys* arose in result of fusions of all ancestral chromosomes into one large chromosome with following extensive rearrangements in its regions. In *D. pachys,* unusual for parthenogenetic species high heterozygosity level is a result of independent evolution of chromosome copies that were genetically isolated through inversions by reducing crossing-over within the inverted regions (Fradin et al., 2017). In contrast to *D. pachys*, the *M. lignano* genome consists of two parts, represented by large and small chromosomes, respectively. Small metacentrics showed high homozygosity, typical for inbred line, while the largest chromosome, which look like a pair of homologs and contains alsmost the entire ancestral genome, showed a high heterozygosity level.

It remains a mystery why chromosome fusions followed by rearrangements and probably genetic isolation of large chromosome copies were limited to the part of the duplicated genome. Accurate assembling *M. lignano* genome and reconstruction of its evolutionary reorganization may provide the answer to this question. Interestingly, the independent WGD events occurred in both the *M. lignano*/*M. janickei* and *M. mirumnovem* lineages were similar in some features. Recent WGD followed by chromosomes fusions and formation of large metacentric chromosome. But in contrast to *M. lignano* and *M. janickei*, further reorganization of arose largest chromosome in *M. mirumnovem* was more dramatic and was detected even with molecular cytogenetic methods (Zadesenets et al., 2020, 2021). Furthermore, at least two types of large chromosomes have been described. They differed in size, location of repeat-enriched regions and in the presence of euchromatic regions paralogous to the that of small metacentrics. In contast to *M. mirumnovem*, the difference between the copies of the largest chromosome in *M. lignano* was only uncovered applying the combined approach using whole-genome sequencing and cytogenetic techniques.

Difference between copies of the largest chromosome of *M. lignano* additionally to their genetic isolation could provide errors in meiosis producing gametes with an extra or missing chromosomes in specimens with the 2n=8 karyotype. High frequency of aneuploids on the chromosome MLI1 in the DV1 line may be considered as indirect evidence for that. Given the genome organization in *M. lignano*, we could expect that doubling of the large chromosome should partially solve meiotic problems mentioned above. There are at least two facts that are in concordance with this sugegstion: (1) the laboratory line with the doubled copy number of the MLI1 is rather stable; (2) *M. janickei* probably originated from *M. lignano* via doubling copy number of its largest chromosome.

The discovery of the unusual structure of the *M. lignano* genome prompted us to look for possible evolutionary scenarios that led to the described organization of the genome after WGD event. Cytogenetic rediploidization of the *M. lignano* genome included fusions of ancestral chromosomes into the large one has accompanied by intensive chromosome rearrangements limited with the arose large chromosome. The latter facilitated chromosome copy differentiation and increasing genomic diversity. Natural selection favoring a pair of differing copies could fix them in a population, especially when it prones to founder events or population bottlenecks. It is noteworthy that different pairs of chromosome copies could be fixed in different populations and, in addition, more than two chromosome variants could be present in the same population. Different chromosome rearrangements in copies of the large chromosome and their genetic isolation led to their further independent evolution and increased genetic diversity, while the remaining, more stable part of genome provided the basic genome functioning essential for survival.

Analyzing the genome sequence of *M. lignano*, we should take into account that the size of the existed haploid genome assembly is about 1.5 times larger than the expected one, based on the estimated genome size, and that specimens from different populations may show prominent differences due to the fixation of different copies of the large chromosome. In result, any attempts to obtain genome assembly of the “expected” size fitted to a tetraploid genome have not been successful due to the specific organization of the *M. lignano* genome.

Our knowledge about the genome heterozygosity level, as well as possible differences between homologous, but heterozygous large chromosome copies in karyotypes of worms from natural populations are limited. Are they characterized with an especial pair of heterozygous copies of the MLI1 or different variants of the MLI1 co-exist in natural populations and fixation of any pair of its copies takes place due to the population bottleneck or inbreeding effect in the laboratory line(s)? Earlier, we showed that karyotype diversity associated with aneuploidies on the largest chromosome took place in natural populations of *M. lignano* and its relative species *M. janickei*. Given the cytogenetic data on their karytoype organization, we inferred the new uncommon mechanism of speciation through divergence and genetic isolation of the large chomosome copies followed by their doubling and fixation of new chromosome variants.

## Conclusion

The consequences of WGD are complex and variable, and seem to differ between different species. In this study, we reported unusual subgenomic composition of the neoautopolyploid species *M. lignano* through the combined application of whole-genome sequencing and molecular cytogenetic techniques, including chromosome microdissection. Here we presented the uncommon mechanism of rediplodization of the polyploid genome *M. lignano*, which consists in (1) presence of three subgenomes, emerged via formation of large fused chromosome and its variants, and (2) sustaining their heterozygosity through inter- and intrachromosomal rearrangements. These results expand our understanding of possible unusual trajectories of karyotype and genome evolutionary reorganization after a recent WGD in animals. Additionally, our data reflect the challenges scientists face during reconstruction of genome having a hidden polyploid origin.

## Materials and methods

### 1. Palatino Linotype> Worm samples

The inbred line of *M. lignano*, DV1, has been established via full-sib and half-sib inbreeding for 24 generations (Janicke et al., 2013) and then was maintained under standard laboratory conditions (Rieger et al., 1988; Ladurner et al., 2005) at small population sizes for a high level of genetic homozygosity. For the last years the inbred DV1 line was regularly karyotyped and a high karyotype instability associated with the variable copy number of the largest chromosomes was revealed (Zadesenets et al., 2016, 2017a, b, Zadesenets and Rubtsov, 2021).

### 2. Palatino Linotype> Experimental design

#### 2.1 Establishing the DV1_8 and DV1_10 sublines, derivatives of the DV1 line

Given the previously uncovered karyotype diversity within the DV1 line (Zadesenets et al., 2016, 2017a, b), we aimed to establish purebred sublines with two different karyotypes, namely: (1) the species-specific 2n=8 karyotype of *M. lignano;* (2) the 2n=10 karyotype with four copies of the largest chromosome. The obtained sublines should be characterized with stable karyotype. We will remind that the established earlier subline DV1/10 (2n=10) has been maintained for four years. Its regular karyotyping, done at different subline ages, revealed from 1 to 5% of specimens showing the karyotypes distinct from the expected one (Zadesenets et al., 2017a, b). Here we established two new sublines, DV1_8 and DV1_10, with the karyotypes 2n=8 and 2n=10, respectively, differing with the copy number of the largest chromosome MLI1 (Fig. S1). The DV1_8 subline showed the species-specific karyotype, two large and six small metacentrics, while DV1_10 showed tetrasomy on the MLI1. Both sublines, DV1_8 and DV1_10 were established simultaneously. Each subline was initiated from two individuals chosen from 50 previously karyotyped individuals of the inbred line DV1. Before the experiment, the DV1_8 and DV1_10 sublines were cultivated for six months (∼8 generations) under standard laboratory conditions (Fig. 4).

**Figure 4.**
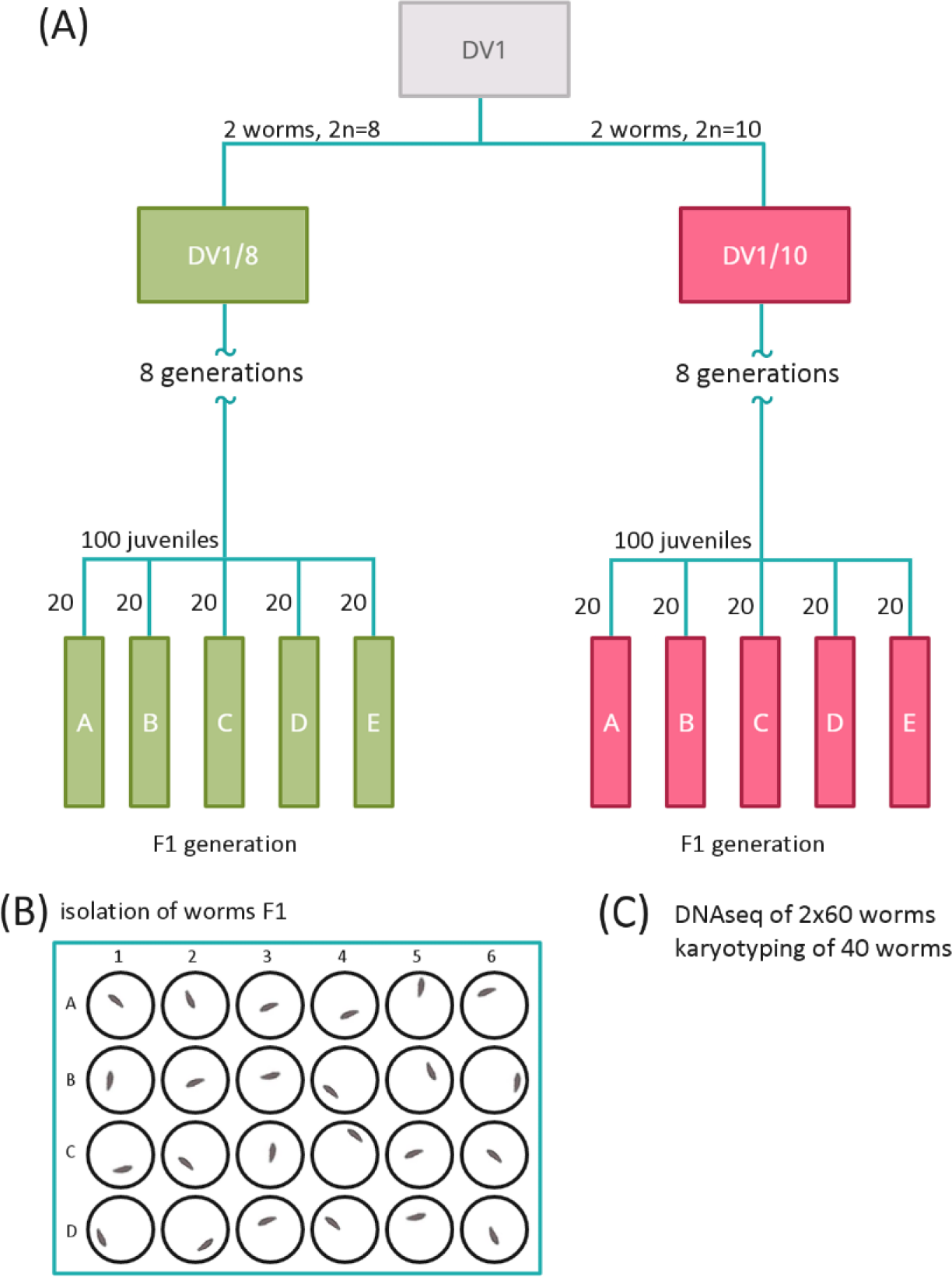
Design of the experiment conducted for this study. (A) Establishment of the sublines DV1_8 and DV1_10 and further generation of worm pools DV1_8A-E and DV1_10A-E. (B) The hatched worms F1 in pools were isolated separately in 24-well plates. (C) Obtaining two replicates of 60 matured F1 worms of the same age from each pool for whole-genome sequencing, in addition, karyotyping of 40 F1 worms from each pool.

#### 2.2 Generation of worm pools from from the sublines DV1_8 and DV1_10

We generated five pools of worms from each subline, DV1_8A-E and DV1_10A-E; each pool was initiated from 20 juvenile (virgin) worms taken from the correspondent subline (Fig. 4; Table S3). After the isolated worms started to produce offspring F1, we isolated the hatchlings separately into 24-well plates until their maturation. Over the next four weeks, the hatched worms were transferred every 2-3 days into the fresh f\2 medium with algae. For collection of the worms of the same age they were fixed in RNAlater (Qiagen, Germany) in four weeks after hatchling. Before fixation, worms were starved for two days. In four weeks, we collected the required number of F1 worms for karyotyping and whole-genome sequencing. For genome sequencing, 120 worms were collected from each pool: the fixed worms were combined in two groups consisted of 60 specimens. In addition, each pool was characterized by karyotyping of 40 specimens. To assess the reproduction level, we counted the number of all hatched worms, dead worms, and odd (morphology and/or behavior) worms in F1 after their maturation. In addition, we karyotyped all worm-founders that were used for generation of the worm pools DV1_8A-E and DV1_10A-E. The pool DV1_8B, in which we detected the specimen with the chromosomal abnormalities, was excluded from further analyses. Finally, we chose three best pools for each subline, DV1_8 and DV1_10 (DV1_8A, DV1_8C, DV1_8E and DV1_10A, DV1_10C, DV1_10E, respectively) according the data on reproduction, mortality, morphology and/or behavior abnormality. We generated two replicate libraries for whole-genome sequencing from each of the selected pool; each library was generated from 60 worms.

### 3. Palatino Linotype> Genomic DNA extraction and preparation of DNA libraries for next-generation sequencing

The pooled worms were kept at -20 °C then gDNA extraction using the standard phenol-chloroform method was performed. For fragmentation, 100 ng of gDNA was incubated with dsDNA Fragmentase (New England Biolabs, USA) at 37 °C for 35 minutes. Libraries of the fragmented DNA were prepared with the NEBNext® Ultra™ II DNA Library Prep Kit for Illumina (New England Biolabs, USA) according to the manufacturer’s protocol.

### 4. Palatino Linotype> NGS sequencing of the DNA libraries DV1_8A, C, E and DV1_10A, C, E

The quality of the generated DNA libraries was assessed on an Agilent 2200 TapeStation (Agilent Technologies, Santa Clara, CA, USA). The quantitative analysis was done using qPCR. To examine the equimolar pooling of the libraries and their quality, a pilot sequencing was done on Illumina MiSeq Nano platform (Illumina, San Diego, CA, USA). The libraries were sequenced on an Illumina NovaSeq 6000 platform (S Prime flow cell) with 2 × 150 bp paired-end reads (Evrogen Joint Stock Company, Russia).

### 5. Palatino Linotype> Generation and sequencing of microdissected DNA library derived from the chromosome MLI1

We generated the microdissected DNA library from 15 copies of the chromosome MLI1 (DV1 line) as was described previously (Zadesenets et al., 2016, 2017a). The initial DNA amplification of the collected chromosomes was performed using a GenomePlex Single Cell Whole Genome Amplification Kit (WGA4) (Sigma-Aldrich, USA) according to the manufacturer’s protocol. Preparing the MLI1 DNA library was performed using NEBNext Ultra II DNA Library Prep kit (New England Biolabs, USA) and sequencing on an Illumina MiSeq with paired-end 300 bp reads at the “Molecular and Cellular Biology” core facility of the IMCB SB RAS (Novosibirsk, Russia).

### 6. Palatino Linotype> Computational analysis

The reference-based analyses were performed using the draft genome assembly of *M. lignano* (Mlig_3_7, GenBank accession: GCA_002269645.1). Most computations were performed in the R environment using basic functions and GenomicRanges package (Lawrence et al., 2013). Repeatitive DNA sequences in the Mlig_3_7 assembly were annotated using RepeatModeler v4.0.7 and RepeatMasker v1.0.8 software (Smit et al., 2015). The details of the conducted analyses are described below.

#### 6.1 Copy-number analysis of the genome assembly Mlig_3_7

Pairwise local self-alignments of the Mlig_3_7 assembly were computed using lastz v1.04.00 (parameters: --ambiguous=n --notransition --step=5 --gap=600,150 --hspthresh=4500 -- ydrop=15000; https://github.com/lastz/lastz). Subsequent chaining and filtering of alignments was done using UCSC chain/net pipeline (axtChain, chainPreNet, chainToPsl, and pslRecalcMatch; http://hgdownload.soe.ucsc.edu/admin/exe/). After removal of false zero-length indels, any overlapping alignments were split at the boundaries of their overlap (Fig. S1). As a result, each fragment of the reference assembly had only full-length alignments of all its homologs, so that its copy number was defined unambiguously. Such groups of homologous fragments were designated as paralogous blocks (PBs). For each PB, the number of SNPs, indels and Levenshtein edit distance were collected (Fig. 1).

Additionally, the available short-read library of the gDNA of *M.lignano* (NCBI SRA: SRR2064573) was mapped to the Mlig_3_7 assembly using bowtie2 v2.3.3 (Langmead et al., 2012). Read counts were computed for each interval within each PB using featureCounts tool (Liao et al., 2014; parameters: -O -M -p -C) and normalized on interval length (reads per kilobase, RPK). Finally, the coverage of PB was calculated as a sum of RPKs of all its members.

#### 6.2 Analysis of 2D k-mer spectra for the DV1_8 and DV1_10 libraries

Raw reads were preprocessed by cutadapt tool to remove low-quality and adapter sequences. All replicates from the DV1_10 group and five replicates from the DV1_8 group were merged into two pools (DV1_8A library was excluded because of substantial GC bias). KMC tools v3.0.0 (Kokot et al., 2017) were used to perform initial k-mer counting (k=35) in two merged datasets and all subsequent logical operations with the resulting k-mer sets (intersect, subtract, subset, dump). Only first reads in a pair were used in k-mer counting to avoid double counting of overlaps in paired reads. Only k-mers ocсurring at least twice in both DV1_8 and DV1_10 sets were kept (99% of all k-mers), which greatly reduced the fraction of erroneous k-mers. Since each k-mer was counted in two datasets, the final frequency spectrum was two-dimensional (DV1_8 and DV1_10, Fig. 2). Interestingly, we observed strong dependence of k-mer frequency distribution on GC-content of k-mers, with GC-rich k-mers provided much higher resolution of the peaks (Fig. S3).

While the bivariate mixed negative-binomial regression seemed to be the most appropriate model for the resulting 2D spectra, we failed to find an affordable ready-to-use solution for this task. Thus, we utilized an approach based on GenomeScope2.0 polyploid models (Ranallo-Benavidez et al., 2020) to infer heterozygosity rates from a pair of marginal spectra (DV1_8 and DV1_10). Innovative approach including generation of karyotype-specific sublines in combination with adapted multivariate k-mer analysis was applied to further analysis.The first spectrum was modeled using ‘predict4_0’ tetraploid model, while the second one - using ‘predict6_0’ hexaploid model. Since at most four allelic states can simultaneously exist in both hexaploid and tetraploid states of the studied genome, we omitted two heterozygosity parameters, ‘raabcde’ and ‘rabcdef’, that describe five or more alternative alleles in the ‘predict6_0’ model. To make the joint fit of two models, we redefined the parameters of ‘predict4_0’ tetraploid model as corresponding combinations of “hexaploid” parameters (Table S1). In order to describe partial correspondences (q_1_ and q_2_), the parameters of proportions, ε_1_ and ε_2_, were introduced. The final nonlinear model formula took the form: ‘Y ∼ length * cbind(X*predict4_0(r1 + eps1*r2 +eps2*r3, (1-eps1)*r2 + (1-eps2)*r3, r4+r5+r6, r7+r8, k, d, kmercov*N, bias*N, X), X*predict6_0(r1, r2, r3, r4, r5, r6, r7, r8, 0, 0, k, d1, kmercov, bias, X))’, where Y is a two-column matrix of transformed marginal k-mer spectra, and N is the normalisation factor for both the sequencing depth (ratio of total k-mer counts) and difference in genome lengths (2 : 3 for the DV1_8 : DV1_10 pair).

#### 6.3 Subgenome classification of Mlig_3_7 scaffolds

The first assessed metric for classification was based on genome coverage by paired reads of the DV1_8 and DV1_10 pooled libraries. Preprocessed reads were mapped to the Mlig_3_7 assembly using bowtie2 v2.3.3 and counted across non-overlapping genomic windows of length 5kb using bedtools v2.30.0 (https://github.com/arq5x/bedtools2). The distribution of genome coverage by the DV1_8 library was bimodal (Fig. 3A), where the lower peak corresponded to two haploid subgenomes (L_1_ and L_2_), and the higher peak - to one diploid subgenome (S). Read counts were normalised on sequencing depth of the highest peak using the ‘median of ratios’ algorithm (Anders et al., 2010). The distribution of the log_2_-ratios of coverage with DV1_8 to DV1_10 reads, log_2_(R_RC_), had the same bimodality, and after centering at the mean between the summits of two peaks, the sign of this metric was used to separate S- and L-class windows (Fig. S4).

The second metric was based on 2D spectrum of k-mer frequencies in the DV1_8 versus DV1_10 libraries and, thus, was more tolerant to the reference misassemblies. Extraction of k-mers specific for S or L_1_ and L_2_ subgenomes was performed using KMC tools (Kokot et al., 2017). As illustrated on Figure 3, k-mers occurring in a range of frequencies (28-39 ∩ 16-25) for the DV1_8 versus the DV1_10 sets were assumed specific for subgenome S (Table S2), and the ranges (28-39 ∩ 36-48) and (12-20 ∩ 16-25) were used to extract a subset of L_1_L_2_-specific k-mers. The subsets of k-mers were mapped to the Mlig_3_7 reference genome using bowtie2, requiring only perfect matches. The number of mapped k-mers over 5-kb non-overlapping genomic windows was normalized to the total number of mapped k-mers, and log_2_-ratio of S-specific to L-specific k-mer counts, log_2_(R_KC_), was used for classification of each genomic window into S and L classes (Fig. 3B).

The final classification scheme was as follows. S-class windows required to be in the top 1/3 windows ranked by coverage with S-specific k-mers and have log_2_(R_KC_) ≥ 1 and positive log_2_(R_RC_). L-class windows required to be in the top 2/3 windows ranked by coverage with L-specific k-mers and have log_2_(R_KC_) ≤ -1 and negative log_2_(R_RC_).

#### 6.4 Processing of WGA-MLI1 library

Isothermal whole genome amplification of extremely low amounts of DNA is associated with several kinds of bias, including overamplification, uneven coverage and prevalence of AT-rich sequences. The sequencing data of WGA-MlLI1 library was passed through multi-step trimming to remove illumina adapters, WGA primers and adjacent poly-GT stretches at the remaining read tails, using cutadapt v3.4 (Martin 2011). Short (< 50 bp) and low-complexity reads were filtered out by bbmap:bbduk tool (v38.91; https://sourceforge.net/projects/bbmap/). The remaining reads were multiply mapped with bbmap:bbmapskimmer (maxsites=40 maxsites2=500 sssr=0.85 secondary=t). Alignment were considered unique if the read is mapped one or more times (at most 10 times, tolerating 1bp mismatch/indel) specifically to only one scaffold in the assembly {get_multimap_info_bbmap.pl}. Breadth of genome coverage in 5-kb genomic windows was calculated and normalised on the number of bases with mappability value of 1 in the window. Here, genome mappability was computed using genmap v1.3.0 tool (-E 2 -K 100 --exclude-pseudo) in scaffold-wise mode. That is, mappability of 100-mer is 1 if it occurs one or more times only in a given scaffold. The resulting fraction of scaffold-specific positions covered by WGA-MLI1 was square root-transformed to enhance the discrimination of near-zero values (WGA-MLI1 score, Fig. 3C).

## Data Access

The sequencing data generated in this study have been submitted to the NCBI BioProject database (https://www.ncbi.nlm.nih.gov/bioproject/) under accession number PRJNA951191.

## Author Contributions

K.S.Z. conceived and designed the experiments, established sublines and worm pools, generated replicates and fixed the material. N.P.B. extracted the genomic DNA and prepared the DNA libraries for sequencing. N.I.E. conducted bioinformatic analysis. K.S.Z., N.I.E., N.B.R. analysed and interpreted the obtained results. N.B.R. obtained funding. K.S.Z., N.I.E., N.B.R. drafted and revised the manuscript. All authors have read and approved the final version of the manuscript.

## Acknowledgements

The authors gratefully acknowledge the resources provided by the “Molecular and Cellular Biology” core facility of the IMCB SB RAS (Novosibirsk, Russia). Microscopy was performed at the Interinstitutional Shared Center for Microscopic Analysis of Biological Objects (ICG SB RAS, Novosibirsk). Resource-intensive calculations were performed at the CCU “Bioinformatics” SB RAS. Generation and sequencing of the gDNA libraries was funded by the Russian Science Foundation (RSF) under grant 19-14-00211. Maintenance of the laboratory lines of *M. lignano*, including the establishment of sublines and worm pools for the current study, was supported by project FWNR-2022-0015.

